# An information thermodynamic approach quantifying MAPK-related signaling cascades by average entropy production rate

**DOI:** 10.1101/431676

**Authors:** Tatsuaki Tsuruyama

## Abstract

Information thermodynamics has recently greatly developed the application for analysis of biological phenomenon. During the signal transduction, entropy production from phosphorylation of signal molecule is produced at individual step production. Using this value, average entropy production rate (AEPR) is computable.

In the current study, AEPR in each signal step was analyzed using experimental data from previously reported studies of the mitogen-activated protein kinases (MAPK) cascade. The result revealed that the differences of AEPR is smaller when using ligands, suggesting that AEPR is one of the attributes of the given cascade and useful for quantitative analysis. This consistency of AEPR suggests that the number of signal events is maximized, in other words, signaling efficiency is maximized. In conclusion, the current information theoretical approach provides not only a quantitative means for comparison of responses to a specified extracellular stimulation, but also a method for evaluation of active cascades.

**Synopsis:** A variety of methods for quantifying intracellular signal transduction have been proposed. Herein, a novel method of quantification by integrated analysis consisting of kinetics, non-equilibrium thermodynamics, fluctuation theorem and graph theory was attempted.

- Signal transduction can be computed by entropy production amount from the fluctuation in the phosphorylation reaction of signaling molecules.
- By Bayesian analysis of the entropy production rates of individual steps, they are consistent through the signal cascade.

## INTRODUCTION

Cell signal cascade network is one of the complicated systems that involve numerous responses to the extracellular stimuli. For example, ligand stimulation simultaneously induces the activation of a variety of enzymatic reactions (Imhof, 2016); thus, multiple pathways are involved. Systems biology has been developed as a theoretical framework that illustrates a global map of signal interactions (Cheong et al, 2011; Govern & ten Wolde, 2014; Selimkhanov et al, 2014; Wang et al, 2010). This framework serves as a statistical model of signaling and gene expression pathways and protein–protein interactions (Drozdov et al, 2013; Edwards et al, 2012; Guo et al, 2017; Sato et al, 2017; Teschendorff et al, 2015; Teschendorff et al, 2017; Teschendorff & Enver, 2017; Teschendorff et al, 2014; White et al, 2006).

In recent decades, information thermodynamics has also greatly developed, and the application of this theory to the analysis of signal transduction is attempted here (Sagawa et al, 2014). Uda et al. have conducted quantitative analysis in the mitogen-activated protein kinases (MAPK) cascade, analyzing robustness from perspectives of mutual entropy (Uda & Kuroda, 2016; Uda et al, 2013). In addition, recent experimental data on signal transduction appears to make it possible to analyze it quantitatively (Kim et al, 2003; Lee et al, 2013; Mackeigan et al, 2005; Mina et al, 2015; Newman et al, 2004; Petropavlovskaia et al, 2012; Tao et al, 2010; Tsuruyama et al, 2016; Tsuruyama et al, 2011; Tsuruyama et al, 2010; Tsuruyama et al, 2002; Wang et al, 2002; Wang et al, 2014; Yeung et al, 1999; Zhang et al, 2011). In earlier studies, we reported that the entropy production rate is one of the parameters that can determine the amount of signal transduction (Tsuruyama, 2014; Tsuruyama, 2018b; Tsuruyama, 2018c; Tsuruyama, 2018d). In these works, in an ideal model of MAPK cascade, the entropy production rate is consistent in the signal transduction step through the cascade. In the current study, we use information thermodynamics (Sagawa et al, 2014) based on the entropy production rate and aim to quantitatively evaluate the entropy production rate using actual experimental data.

## RESULTS

### A model cascade

The modeled cascade is given as a sequential chemical reaction. The signaling molecule at step *j* of cascade *m* is denoted as *X_mj_*, which induces the phosphorylation of *X_mj+1_* into *X_mj+1_*\*. In individual steps, the dephosphorylation of *X_mj+1_*\* into *X_mj+1_* occurs spontaneously or via an enzymatic reaction catalyzed by the *j* phosphatase (*D_mj_*; 1 ≤ *j* ≤ *n*). This reversible signaling step in the above cascades may be described as follows:

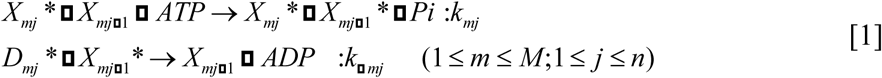

where *k_mj_* and *k_-mj_* are the kinetic coefficients, and ATP, ADP, and Pi represent adenosine triphosphate, adenosine diphosphate, and inorganic phosphate, respectively. When the ligand stimulates the cell, the phosphorylated signaling molecules *X_mj_* is tentatively increased for *τ_+mj_* (> 0), followed by a decline over a long duration, *τ_-mj_* (< 0) (Fig. 1). For example, signaling cascades have been studied extensively using data of mitogen-activated protein kinase (MAPK) pathways, in which the epidermal growth factor receptor (EGFR), *c*-Raf, MAP kinase-extracellular signal-regulated kinase(MEK), and kinase-extracellular signal-regulated kinase (ERK) are sequentially phosphorylated after stimulation with epidermal growth factor (EGF) as follows:

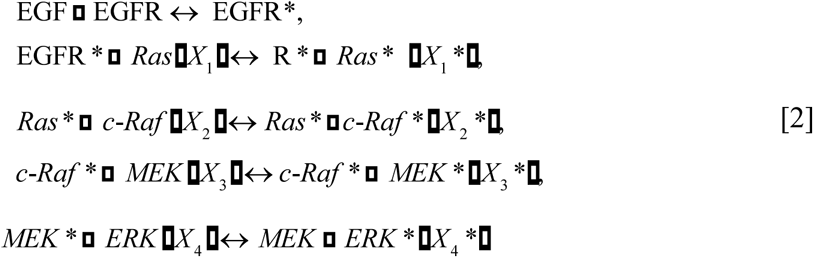

**Fig. 1.**
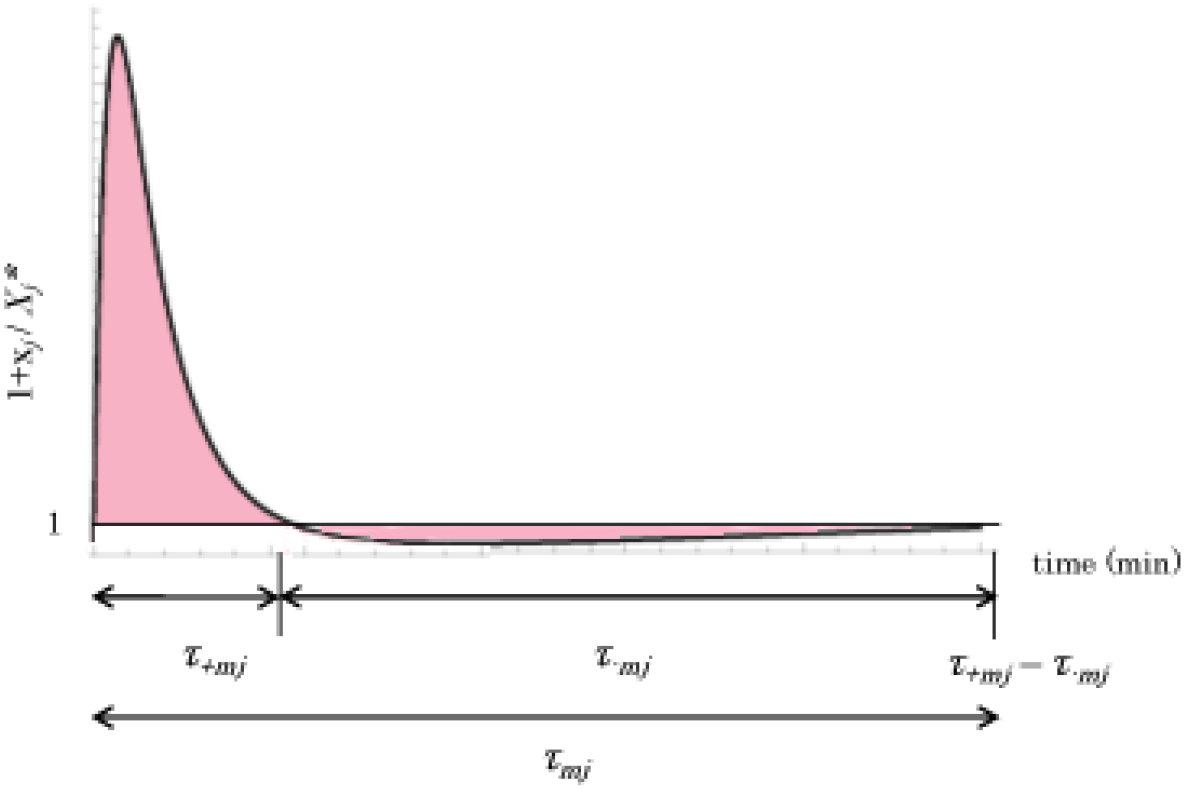
Common time course of step *j*. Fold changes in phosphorylation are shown. The vertical axis denotes the ratio (*X_mj_*\*^st^ +*x_mj_*\*)/*X_mj_*\*^st^. The horizontal axis denotes the duration (min) of step *j. τ_mj,_* represents the total duration of step *j*. A common time course of the *j*^th^ step. Fold changes in modification. *τ_mj_* and *τ-_mj_* represent the duration of the *j*^th^ step and the –*j*^th^ step, respectively. The line ratio = 1 denotes the ratio at the steady state. The integral area is indicated by red colored area.

### Average entropy production rate in MAPK models

At first, AEPR <*σ_mj_*> was introduced for duration *τ_+mj_* – *τ-_mj_* of *j* step using the entropy production rate *σ_mj_* (*t*) that depends on time passage of the step in the signal cascade:

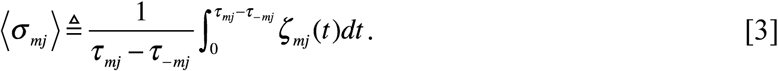

The positive and negative values were assigned to *τ_+mj_* and τ_*-mj*_ by consideration of orientation of the signal cascade. Here, *t* is an arbitrary parameter representing the progress time of a signal event. When the cell signaling system is not far from equilibrium state at *detailed balance* around the *homeostasis*, as follows:

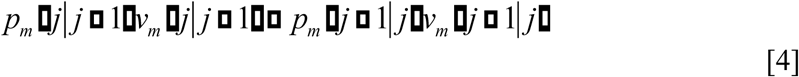

In above, *v* (*j*+1|*j*) and *v* (*j*| *j*+1) signify the transitional rate from the step *j* to the step *j*+1 and the transitional rate from the step *j*+1 to *j* step, respectively. *p* (*j* + 1|*j*) and *p* (*j* |*j* + 1) signify the transitional probability of step *j+*1 given step *j*, and the transitional probability of step *j* given *j* + 1 step, respectively. According to [4], the transitional rate from the step *j* to the step *j*+1, *v* (*j*+1|*j*), is obtained by using the kinetic coefficient, *c_j_*, for step *j*:

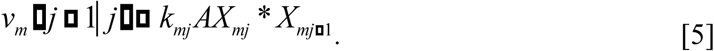

Furthermore, the transitional rate from the step *j*+1 to *j* step, *v* (*j* |*j*+1), which is equal to the de-modification reaction of the backward signal transduction, is given by

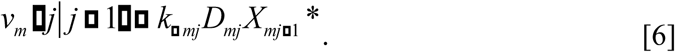

We investigated the relationship between chemical work and information based upon a model in [1] or [2]. For this purpose, the average entropy production rate of an actual fluctuating signaling cascade is quantified.

From Eqs [4], [5], and [6], the following is obtained:

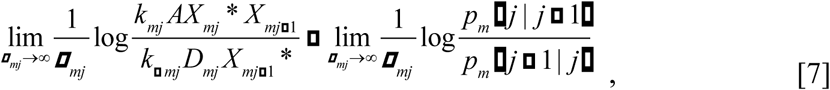

and the fluctuation theorem (FT) gives:

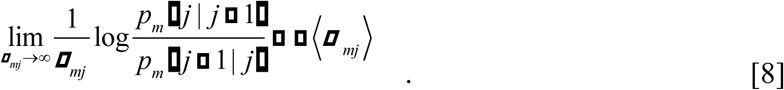

From [7] and [8], we have

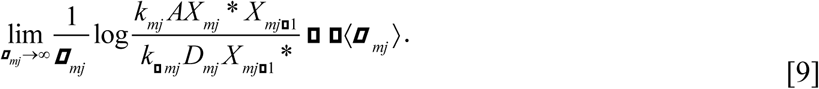

Here, let us consider the fluctuation from the steady state. From the concentration of the signaling molecules at the homeostatic state using the superscript *st* and the concentration fluctuation, the following is obtained:

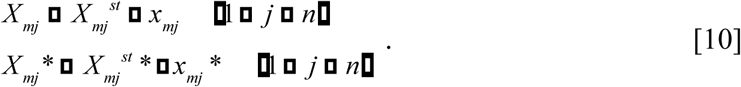

Using the fluctuation of the signaling molecule *x_mj_* and the concentration at the equilibrium state 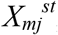, we obtain the following from [9] and [10]:

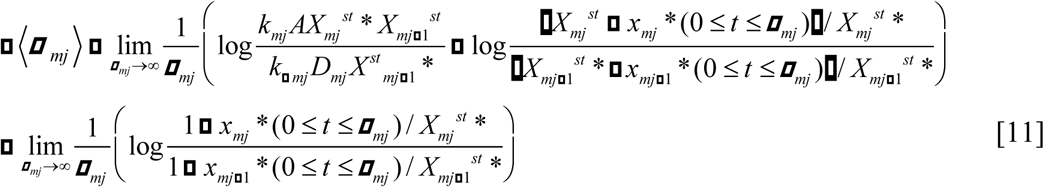

Considering that *x_mj_*\* can be regarded significantly smaller than *x_mj+1_*\* during the signal step *j* according to experimental data (Kim et al, 2003; Lee et al, 2013; Mackeigan et al, 2005; Mina et al, 2015; Newman et al, 2004; Petropavlovskaia et al, 2012; Tao et al, 2010; Tsuruyama et al, 2016; Tsuruyama et al, 2011; Tsuruyama et al, 2010; Tsuruyama et al, 2002; Wang et al, 2002; Wang et al, 2014; Yeung et al, 1999; Zhang et al, 2011), <*σ_mj_*> is approximately given by:

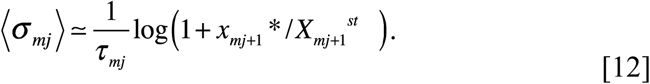

Eq. [11] implies that the duration *τ_mj_* is expressed in a linear relationship with the log arithm of the concetration fluctuation. This simple relation may be validated by using experimental data as follows. In earlier studies, we deduced that the entropy production rate <*σ_mj_*> is independent of the step number *j* (Tsuruyama, 2018a):

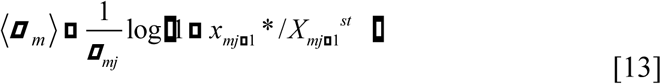

Below, we discuss whether AEPR is independent of *j*.

### Bayesian statistical analysis of the AEPR of the Ras-cRaf-ERK cascade

Here, AEPR was calculated using the data of previous experiments (Blossey et al, 2012; Hollenberg, 2002; Lee et al, 2013; Purutçuoğlu & Wit, 2012; Qiao et al, 2007; Xin et al, 2011; Yoon & Deisboeck, 2009; Zumsande & Gross, 2010). In particular, we used the MAPK pathway, in which epidermal growth factor receptor (EGFR/ErbB1 or ErbB2), Ras, c-Raf, MAPK/extracellular signal-regulated kinase (ERK) (Jung et al), and ERK are phosphorylated constitutively following stimulation of the cell with EGF or other agents. The Ras-c-Raf-ERK cascade comprises a ubiquitous signaling pathway that conveys mitogenic and differentiation signals from the cell membrane to the nucleus. Phosphorylation was evaluated by the intensity of the immunoblot of phosphorylated proteins that was normalized by the division by the intensity values in the absence of the ligand stimulation. The data were then analyzed by Bayesian statistics. The ratio

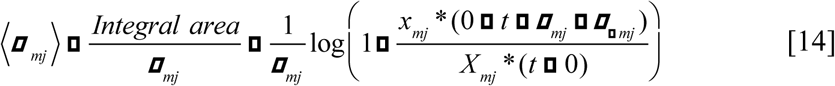

in the right side of Eq. [14] was obtained by measurement of the blot of the protein from the start to the end of the signal event by stimulation. The density of western blot analysis for calculation of the ratio was utilized for computing of AEPR of duringthe phosphorylation of signaling molecules.

### AEPR of MEK-ERK-RSK cascade activation

As an example, we aimed to validate our method using another experimental data through liver *X* receptor activation, which inhibits melanogenesis through the acceleration of ERK-mediated MITF (Microphthalmia-associated transcription factor) degradation (Lee et al, 2013) (Fig. 2). According to this data, AEPR during the phosphorylation is estimated from 0.087 to 0.100 (J/Kmin). Together with above data, we hypothesized that cRaf, MEK, ERK and MSK (mitogen- and stress-activated protein kinase) forms a sequential activation cascade and AEPRs are nearly equal to each other (Table 1). As Wang et al reported (Wang et al, 2014), a set of signal proteins, such as cRaf, MEK, ERK, p38 mitogen-activated protein kinases, JNK(c-Jun NH2-terminal kinases), RSK(ribosomal S6 kinase), and MSK, are considered to participate in the pathway. As a result, several AEPRs during phosphorylation of cRaf, MEK, ERK, MSK were nearly 0.075 (J/Kmin) except RSK (0.086 (J/Kmin)) and p38 (0.096 (J/Kmin)) (Table 2). These data suggest that AEPR is nearly consistent during MEK-ERK-MSK cascade.

**Fig. 2.**
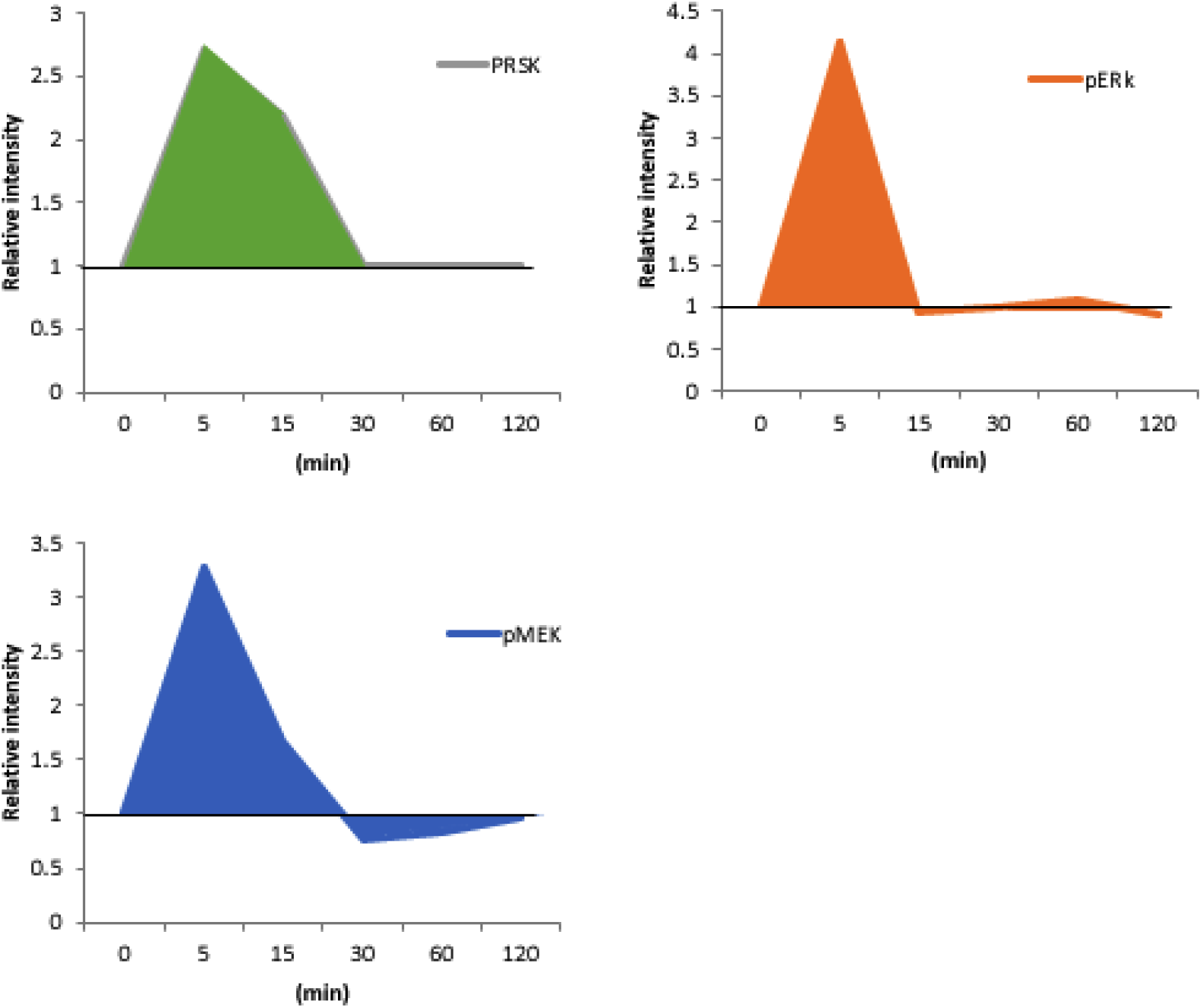
Fold changes in phosphorylation using Liver X. The vertical axis denotes relative intensity ratio of immunoblot signals for *X_mj_*\*. The ratio is computed by (*X_mj_*\*^st^ +*x_mj_\**)/*X_mj_*\*^st^. The horizontal axis denotes the duration (min) of step *j*. The vertical axis label 1 indicates the ratio at the steady state (Lee et al, 2013). The integral areas are indicated by green, orange, and blue colored area.

**Table 1.** Time course of phosphorylation of signaling molecules in MEK-ERK-RSK signaling pathway. The integral areas were calculated by identical method shown in Table 1 legend. A lowercase “p” represents the phosphorylated species. τ, duration of individual steps.

| | 0(min) | 5(min) | 15(min) | 30(min) | 60(min) | 120(min) | $S = (X^{st}+x)/ X^{st}$ | LogS | AEPR |
| --- | --- | --- | --- | --- | --- | --- | --- | --- | --- |
| pMEK | 1.000 | 3.267 | 1.664 | 0.761 | 0.840 | 0.967 | 13.680 | 2.616 | **0.0872** |
| pERK | 1.000 | 3.630 | 1.240 | 0.982 | 1.086 | 0.918 | 20.459 | 3.019 | **0.1006** |
| pRSK | 1.000 | 2.713 | 2.180 | 1.002 | 1.000 | 1.000 | 15.782 | 2.759 | **0.0920** |

**Table 2.**
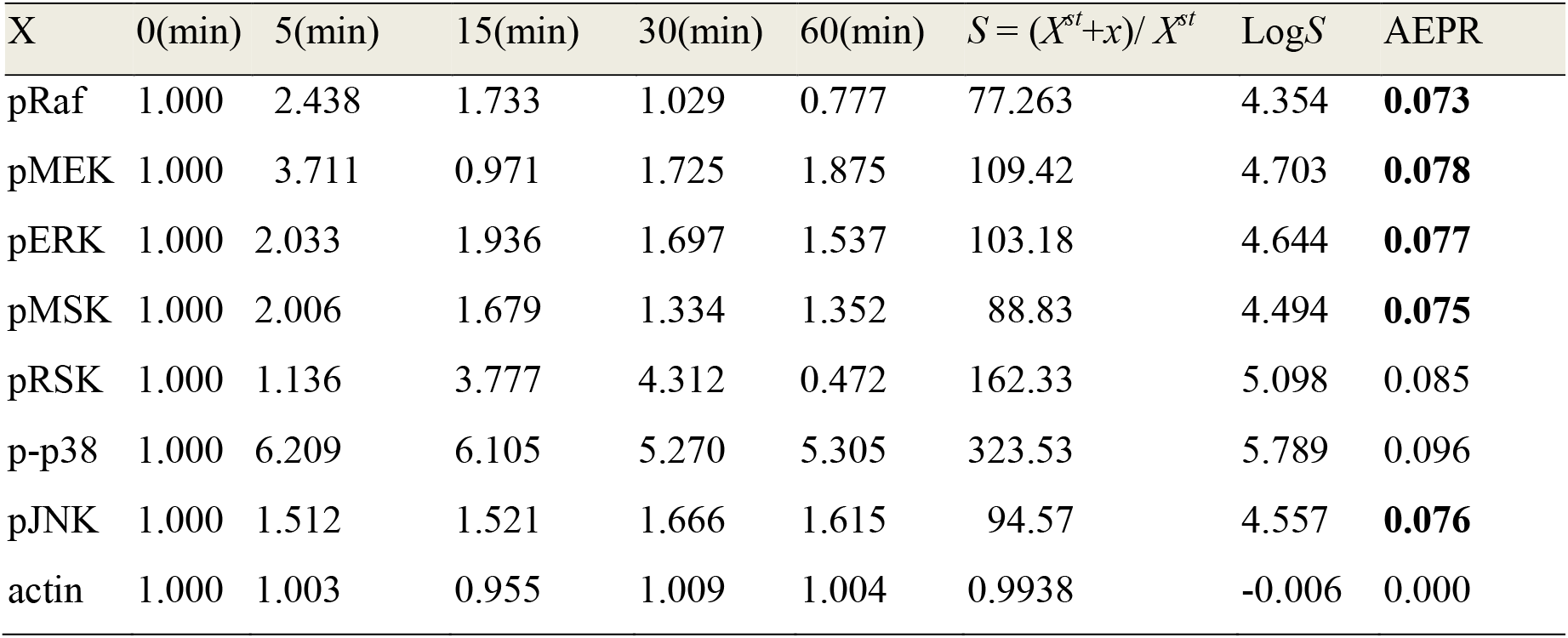
Time course plot of phosphorylation of various signaling molecules in in the signaling pathways. The integral areas were calculated by identical method shown in Table 1 legend. A lowercase “p” represents the phosphorylated species. Minutes represent the measurement time of phosphorylation ratio.

| X | 0(min) | 5(min) | 15(min) | 30(min) | 60(min) | $S = (X^{st}+x)/ X^{st}$ | LogS | AEPR |
| --- | --- | --- | --- | --- | --- | --- | --- | --- |
| pRaf | 1.000 | 2.438 | 1.733 | 1.029 | 0.777 | 77.263 | 4.354 | **0.073** |
| pMEK | 1.000 | 3.711 | 0.971 | 1.725 | 1.875 | 109.42 | 4.703 | **0.078** |
| pERK | 1.000 | 2.033 | 1.936 | 1.697 | 1.537 | 103.18 | 4.644 | **0.077** |
| pMSK | 1.000 | 2.006 | 1.679 | 1.334 | 1.352 | 88.83 | 4.494 | **0.075** |
| pRSK | 1.000 | 1.136 | 3.777 | 4.312 | 0.472 | 162.33 | 5.098 | 0.085 |
| p-p38 | 1.000 | 6.209 | 6.105 | 5.270 | 5.305 | 323.53 | 5.789 | 0.096 |
| pJNK | 1.000 | 1.512 | 1.521 | 1.666 | 1.615 | 94.57 | 4.557 | **0.076** |
| actin | 1.000 | 1.003 | 0.955 | 1.009 | 1.004 | 0.9938 | -0.006 | 0.000 |

As another example, the data of the Ras-cRaf-ERK cascade were used (Petropavlovskaia et al, 2012). These data pertain to the activation of cascades in RIN-m5F rat islet cells stimulated by two types of ligands: full-length recombinant islet neogenesis-associated protein (rINGAP) and a 15-amino acid fragment of INGAP, termed INGAP-P (Table 3). Regarding the data of INGAP-P stimulation, EAP (μ: the Expected A Posteriori) of AEPR was calculated by using Bayesian statistical method for point estimation. The EAPs for the 95% credible interval were 9.90, 9.48, and 8.72 (mJ/Kmin). Meanwhile, the AEPR in the phosphorylation process of AKT was 1.36 (mJ/Kmin), which was significantly less than AEPR in the phosphorylation of Ras, Raf, and ERK. Furthermore, the differences in AEPR between Ras and c-Raf, Ras and ERK, and c-Raf and ERK were 0.421 (mJ/Kmin), –1.579 (mJ/Kmin), and –2.00 (J/Kmin) for the 95% credible interval, respectively (supplementary Table 1; supfig1). From these data, the differences in AEPR in the phosphorylation of Ras, c-Raf, and ERK were smaller, whereas the difference between AEPR of AKT phosphorylation and the AEPRs of Ras, c-Raf, and ERK are –7.101, –6.68 to –8.680 (J/Kmin), which are definitely greater than those of other values of difference among those values of Ras, c-Raf, and ERK. Details of the descriptive statistical values are shown in supplementary Table 1.

**Table 3.**
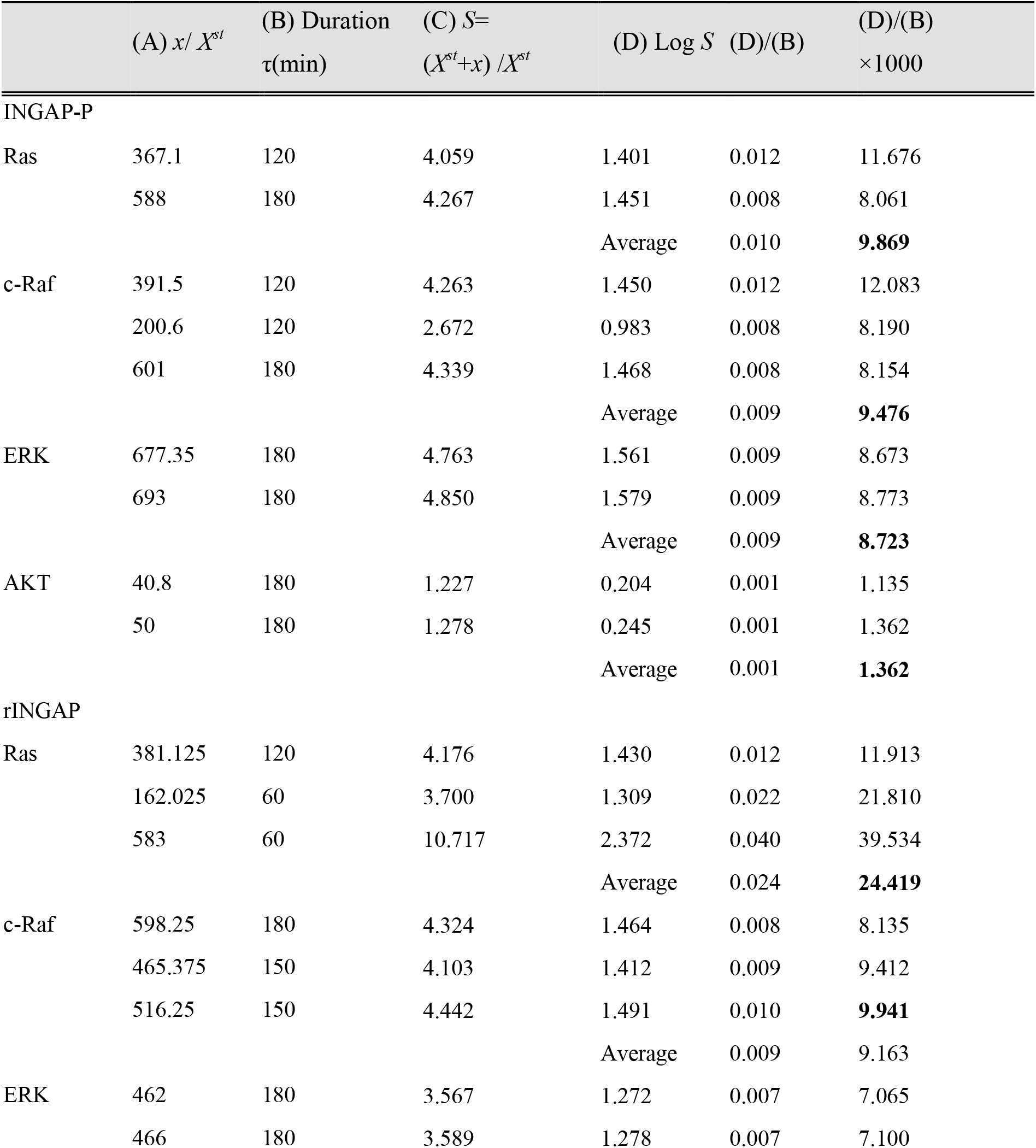
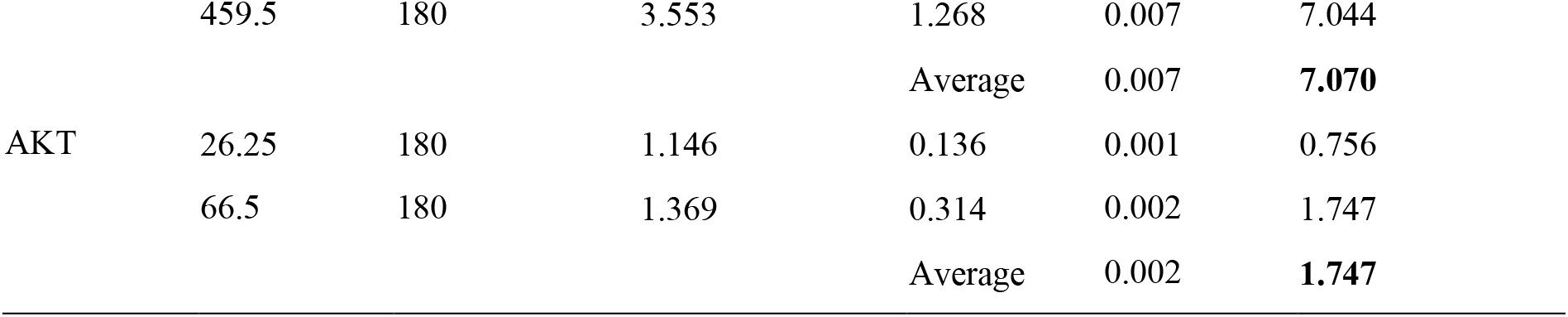
Numerical parameters of the INGAP-P- and rINGAP-induced signaling pathways. The integral areas were calculated by the sum of the fluctuation *S* = 1+ *x*/ *X^s^*^*t*^ from *t* = 0 to *t = τ_m_*_*j*_ (Petropavlovskaia et al, 2012). Bold values were used in the computational calculation (as in the paragraph “*AEPR of MEK-ERK-RSK cascade activation***”** in the results section). Experiments were performed three times. When sufficient activation of cascade was not observed, the data was omitted (phosphorylation of Ras, Erk, AKT in use of INGAP-P; phosphorylation of AKT in use of rINGAP).

Next, the data on rINGAP stimulation were analyzed. The EAPs (μ) of AEPR values in the phosphorylation of Ras, Raf, and ERK were 14.7, 16.4, and 1.68 (mJ/Kmin), respectively (Table 3). Then the differences between the AEPRs of Ras and Raf, Raf and ERK, and Ras and ERK were 14.7, 16.4, and 1.68 (mJ/Kmin), respectively, of which the former two were much larger than those in INGAP-P. On the other hand, the differences of AEPR from the phosphorylation of AKT were –18.97, –4.267, –2.587 (mJ/Kmin), respectively. These data clearly show that the AEPRs of Ras, Raf, and ERK in using rINGAP are significantly variable and might differ from AEPR of AKT (supplementary Table 2; supplementary fig 2). From these data, the activation of the cascade using rINGAP is not consistent, which is distinctly from when using INGAP-P (Fig. 3). Probably, AKT was not involved in this activation cascade Thus, AEPR values may be a promising indicator of signal transduction activity using individual ligands. Petropavlovskaia et al emphasized that INGAP-P has sufficient activity (Petropavlovskaia et al, 2012).

**Fig. 3.**
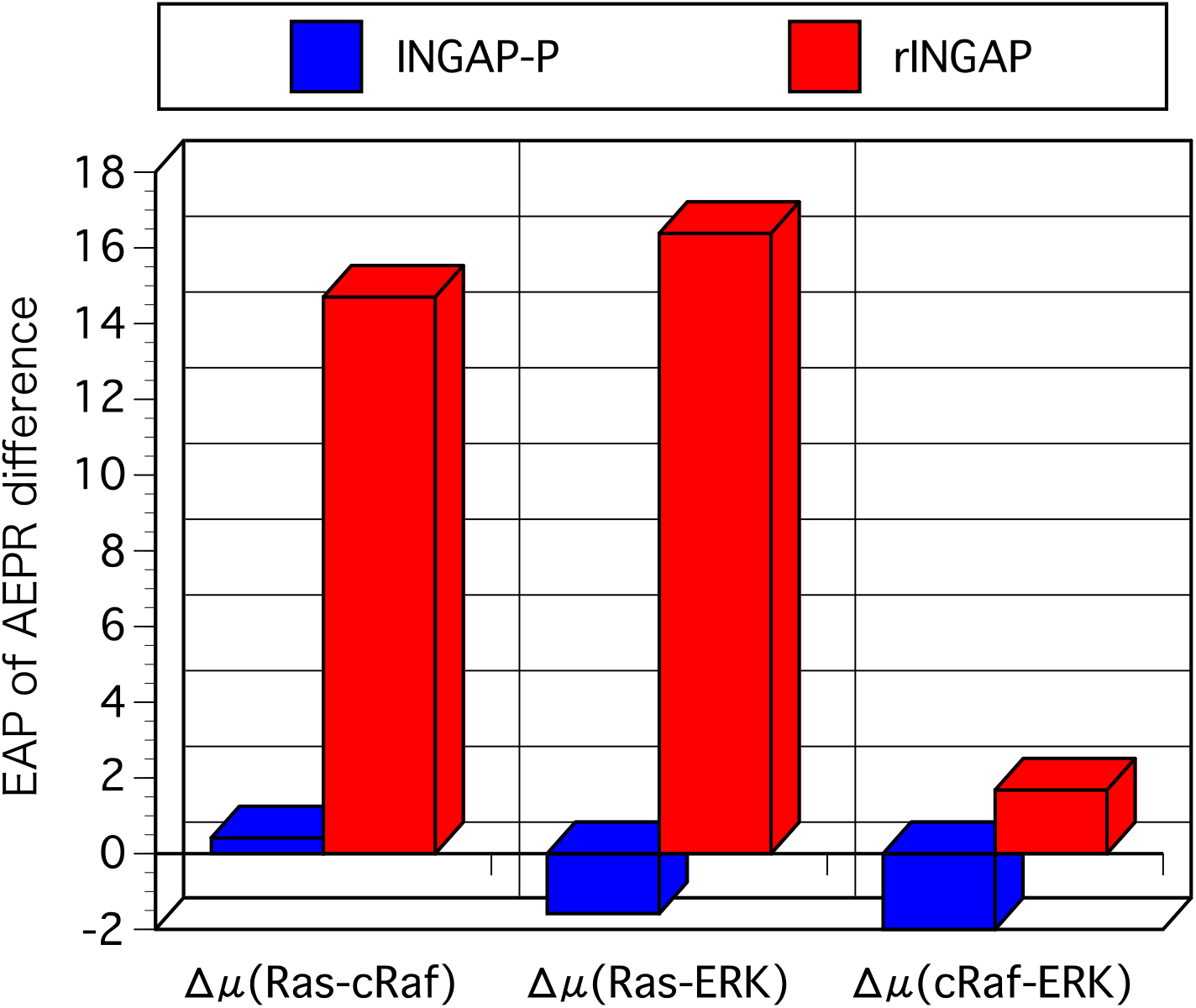
EAP, the expected a posteriori (Δμ: the mean of difference between AEPR during phosphorylation ratio of Ras, cRaf, and ERK in the predictive distribution).

## DISCUSSSION

In the current study, the sequential increase in the concentration fluctuation of phosphorylated signaling molecules showed that the signal transduction along the cascade could be evaluated quantitatively. Quantitative studies of three experiments shown in this study revealed that AERP value is nearly consistent with the MEK-ERK-MSK cascade and MAPK pathway (Lee et al, 2013; Wang et al, 2002) except the data in the stimulation by rINGAP (Petropavlovskaia et al, 2012).

In the following, we discuss quantitative studies of Petropavlovskaia’s experiments in detail (Petropavlovskaia et al, 2012). The differences in the stimulation of INGAP-P and rINGAP-P were investigated using Bayesian statistics for the quantitative evaluation of the AEPR at each step. rINGAP is a full-size protein having normal physiological activity, whereas INGAP-P is an oligopeptide and is a non-physiological substance that is developed for MAPK cascade activation. When using INGAP-P, the differences of AEPR (Δμ) between phosphorylation process of Ras and Raf step, Raf and ERK step, and Ras and ERK steps were sufficiently smaller, suggesting that APER can be considered equal and that this substance induced consistent activation of Ras-cRaf-ERK pathway. In contrast, rINGAP stimulation demonstrated the significant higher value of APER in the phosphorylation process of Ras. This can be interpreted as follows: rINGAP has a more effect on upstream of the signal transduction pathway. On the other hand, whereas difference of mean of AEPR between the phosphorylation process of Raf and ERK was smaller. The consistency of AEPR in using INGAP-P can be related to physiological activity. In previous study, when the signal event number per a given duration is maximized, we obtained the following result (Tsuruyama, 2017; Tsuruyama, 2018c):

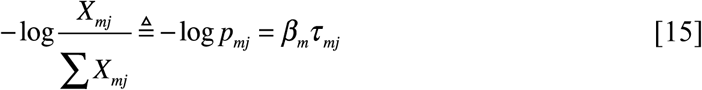

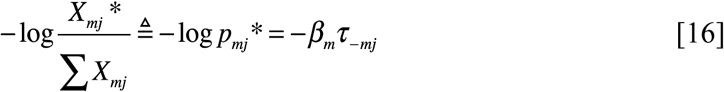

In above, *β_m_* represents an arbitrary constant independent of step number *j*. Above Eqs. [15] and [16] represents a well-known type or entropy coding (Brillouin L, 2013).

According to experimental data (Kim et al, 2003; Lee et al, 2013; Mackeigan et al, 2005; Mina et al, 2015; Newman et al, 2004; Petropavlovskaia et al, 2012; Tao et al, 2010; Tsuruyama et al, 2016; Tsuruyama et al, 2011; Tsuruyama et al, 2010; Tsuruyama et al, 2002; Wang et al, 2002; Wang et al, 2014; Yeung et al, 1999; Zhang et al, 2011), the backward durations were significantly longer.

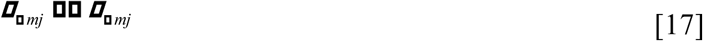

Further, we obtained the following result :

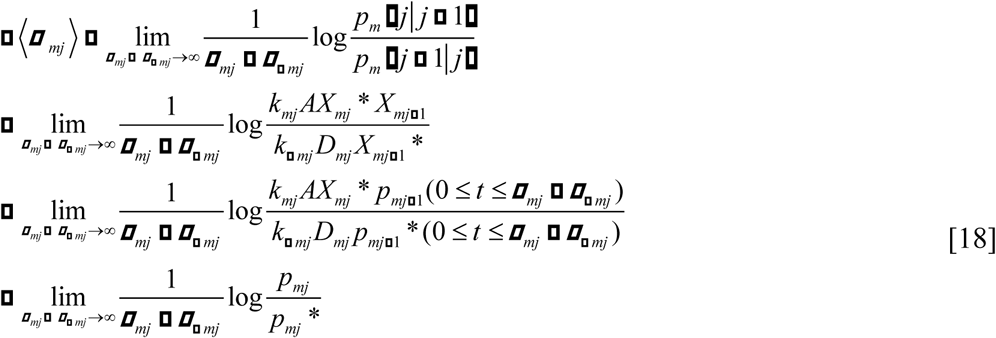

From [15]-[18],

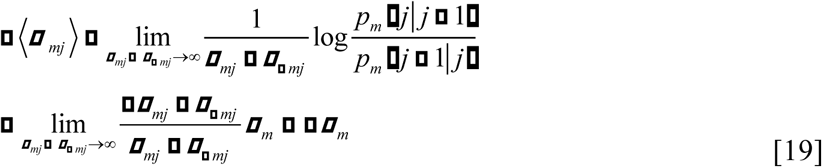

Therefore, AEPR <*σ_mj_*> is equal to *β_m_*, which is independent of the step number *j* (Tsuruyama, 2017; Tsuruyama, 2018a; Tsuruyama, 2018b; Tsuruyama, 2018c).

In the future, further study will be required to prove which strategy of signal cascade the biological system will select. In particular, the cell system may select a strategy to maximize signal event number during a given duration, or accuracy of the signal transduction may be required in the cell system. Cost-performance of metabolomics substances tradeoffs for cellular regulatory functions and information processing has been recently argued in the chemotaxis of *E. coli* (Lan et al, 2012). They pointed out that the framework of information thermodynamics has still been developed. In this way, the strategy chosen for signal cascade by the cell system will likely be determined experimentally. The methodology described herein might provide the quantitative basis for the estimation of ligand or drug effects involving the activation of cellular signal transduction. This theoretical approach appears suitable for the identification of novel active signaling cascades among response cascades.

### Materials and Methods

For the Markov chain Monte Carlo (MCMC) method, the *R-* program (https://www.r-project.org/; Rstan; http://mc-stan.org/users/interfaces/rstan) was used, and a random number was generated to obtain posterior predictive distribution of the average entropy production rate (AEPR). The posterior probability distribution was calculated by the remaining five Markov chains (21,000 times per run) after the removal of the burn-in times. The posterior distribution was approximately predicted by 100000 random numbers obtained by the HMC method. The convergence judgment index 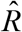 was less than 1.1 and the random number sequence converged to the posterior distribution. From Eq. [11], log (1 + *x_mj_*\*/*X_mj_*\*) of each step is divided by the signal duration *τ_mj_* to obtain the AEPR.

Abbreviation used in the text are PPI, protein–protein interaction); AEPR, average entropy production rate; FT, fluctuation theorem; MAPK, mitogen-activated protein kinase; ERK, extracellular signal-regulated kinase. Densitometric image analysis of previous reported data was performed using Image J software (http://rsb.info.nih.gov/ij/).

## Acknowledgments

We thank Kenichi Yoshikawa of Doshisha University, for his advice.

## SUPPLEMENTARY MATERIALS

### Supplementary figures

**Supplementary figure 1.**
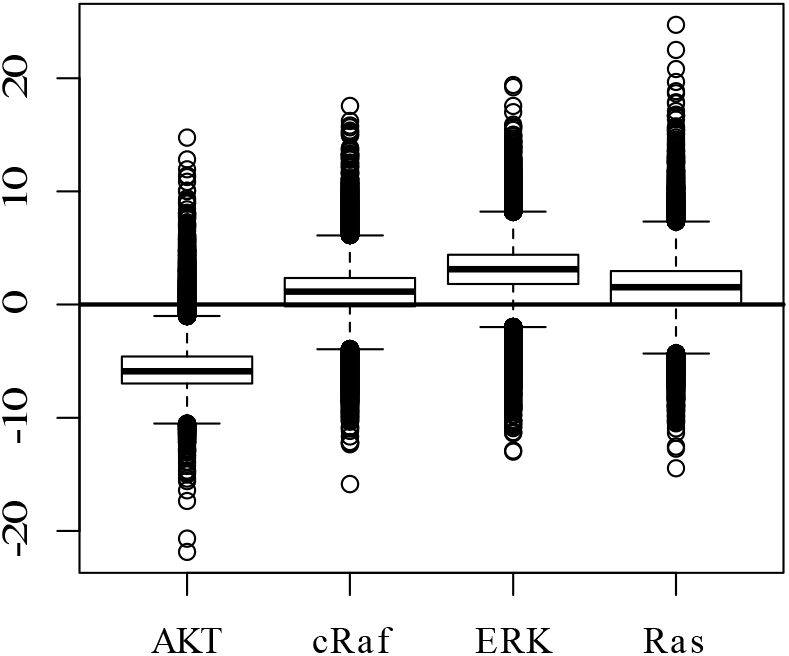
Box-and-whisker plot of posterior distribution of the effect levels (cRaf, ERK, Ras) in INGAP-P stimulation. The vertical axis represents the effect level (difference from the whole average). The vertical axis indicates AEPR (mJ/Kmin).

**Supplementary figure 2.**
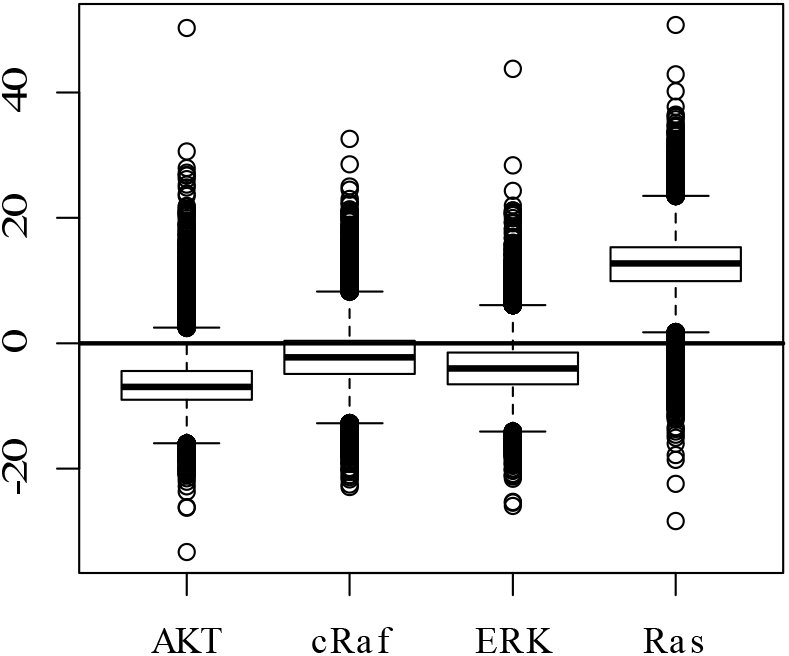
Box-and-whisker plot of posterior distribution of the effect levels (cRaf, ERK, Ras) in rINGAP stimulation. The vertical axis represents the effect level (difference from the whole average). The vertical axis indicates AEPR (mJ/Kmin).

### SUPPLEMENTARY TABLES

**Supplementary table 1.** Bayesian statics for computing level of difference in AEPR Δμ using INGAP-P stimulation during phosphorylation of signal molecules, Ras, cRaf, ERK, and AKT. EAP, expected a posteriori; post.st, posterior sandard deviation; Δμ, difference of mean of AEPR in use of INGAP-P or rINGAP for stimulation. Nonoverlap represents, Cohen’s U3, the third measure of non-overlap; π_d_, represents probability of difference between Xi and Xj; πc, probability beyond threshold. The values in the main text is shown in bold font.

Ras vs. cRaf
|  | EAP | post.sd | 2.50% | 5% | 50% | 95% | 97.50% |
| --- | --- | --- | --- | --- | --- | --- | --- |
| $\Delta\mu$ | **0.421** | 3.847 | -7.294 | -5.744 | 0.408 | 6.623 | 8.235 |
| Size effect | 0.115 | 0.905 | -1.654 | -1.373 | 0.112 | 1.607 | 1.889 |
| Nonoverlap | 0.534 | 0.272 | 0.049 | 0.085 | 0.545 | 0.946 | 0.971 |
| $\pi_d$ | 0.527 | 0.216 | 0.121 | 0.166 | 0.532 | 0.872 | 0.909 |
| $\pi_c$ (0.005) | 0.527 | 0.216 | 0.121 | 0.166 | 0.531 | 0.872 | 0.909 |

Ras vs. ERK
|  | EAP | post.sd | 2.50% | 5% | 50% | 95% | 97.50% |
| --- | --- | --- | --- | --- | --- | --- | --- |
| $\Delta\mu$ | **-1.579** | 3.887 | -9.367 | -7.791 | -1.617 | 4.702 | 6.389 |
| Size effect | -0.442 | 0.921 | -2.245 | -1.962 | -0.443 | 1.077 | 1.365 |
| Nonoverlap | 0.373 | 0.264 | 0.012 | 0.025 | 0.329 | 0.859 | 0.914 |
| $\pi_d$ | 0.397 | 0.213 | 0.056 | 0.083 | 0.377 | 0.777 | 0.833 |
| $\pi_c$ (0.005) | 0.396 | 0.213 | 0.056 | 0.082 | 0.377 | 0.777 | 0.833 |

cRaf vs. ERK
|  | EAP | post.sd | 2.50% | 5% | 50% | 95% | 97.50% |
| --- | --- | --- | --- | --- | --- | --- | --- |
| $\Delta\mu$ | **-2.000** | 3.464 | -8.880 | -7.511 | -2.014 | 3.566 | 5.082 |
| Size effect | -0.557 | 0.830 | -2.190 | -1.921 | -0.555 | 0.808 | 1.065 |
| Nonoverlap | 0.334 | 0.240 | 0.014 | 0.027 | 0.289 | 0.791 | 0.857 |
| $\pi_d$ | 0.367 | 0.193 | 0.061 | 0.087 | 0.347 | 0.716 | 0.774 |
| $\pi_c$ (0.005) | 0.367 | 0.193 | 0.061 | 0.087 | 0.347 | 0.716 | 0.774 |

AKT vs. Ras
|  | EAP | post.sd | 2.50% | 5% | 50% | 95% | 97.50% |
| --- | --- | --- | --- | --- | --- | --- | --- |
| $\Delta\mu$ | **-7.101** | 3.686 | -13.696 | -12.528 | -7.356 | -0.74 | 1.08 |
| Size effect | -2.020 | 1.165 | -4.372 | -3.991 | -1.996 | -0.141 | 0.187 |
| Nonoverlap | 0.093 | 0.153 | 0.000 | 0.000 | 0.023 | 0.444 | 0.574 |
| $\pi d$ | 0.135 | 0.151 | 0.001 | 0.002 | 0.079 | 0.460 | 0.553 |
| $\pi c$ (0.005) | 0.135 | 0.15 | 0.001 | 0.002 | 0.079 | 0.460 | 0.552 |

AKT vs. cRaf
|  | EAP | post.sd | 2.50% | 5% | 50% | 95% | 97.50% |
| --- | --- | --- | --- | --- | --- | --- | --- |
| $\Delta\mu$ | **-6.680** | 3.284 | -12.361 | -11.384 | -6.983 | -0.907 | 0.728 |
| Size effect | -1.906 | 1.071 | -4.062 | -3.705 | -1.886 | -0.174 | 0.126 |
| Nonoverlap | 0.096 | 0.148 | 0.000 | 0.000 | 0.03 | 0.431 | 0.55 |
| $\pi d$ | 0.141 | 0.146 | 0.002 | 0.004 | 0.091 | 0.451 | 0.535 |
| $\pi c$ (0.005) | 0.141 | 0.146 | 0.002 | 0.004 | 0.091 | 0.451 | 0.535 |

AKT vs. ERK
|  | EAP | post.sd | 2.50% | 5% | 50% | 95% | 97.50% |
| --- | --- | --- | --- | --- | --- | --- | --- |
| $\Delta\mu$ | **-8.680** | 3.321 | -14.364 | -13.386 | -8.997 | -2.876 | -1.168 |
| Size effect | -2.463 | 1.208 | -4.911 | -4.498 | -2.433 | -0.531 | -0.193 |
| Nonoverlap | 0.056 | 0.114 | 0.000 | 0.000 | 0.007 | 0.298 | 0.423 |
| $\pi d$ | 0.092 | 0.121 | 0.000 | 0.001 | 0.043 | 0.354 | 0.446 |
| $\pi c$(0.005) | 0.092 | 0.121 | 0.000 | 0.001 | 0.043 | 0.353 | 0.445 |

**Supplementary table 2.** Bayesian statics for computing level of difference in AEPR using rINGAP stimulation during phosphorylation of signal molecules. EAP, expected s posteriori; post.st, posterior sandard deviation; Δμ, difference of mean of AEPR in use of INGAP-P or rINGAP for stimulation. Nonoverlap represents, Cohen’s U3, the third measure of non-overlap; πd, represents probability of difference between Xi and Xj; πc, probability beyond threshold.

Ras vs. cRaf
|  | EAP | post.sd | 2.50% | 5% | 50% | 95% | 97.50% |
| --- | --- | --- | --- | --- | --- | --- | --- |
| $\Delta\mu$ | **14.702** | 7.182 | -0.356 | 2.748 | 14.947 | 25.899 | 28.313 |
| Size effect | 1.835 | 0.967 | -0.031 | 0.261 | 1.824 | 3.445 | 3.758 |
| Nonoverlap | 0.907 | 0.138 | 0.488 | 0.603 | 0.966 | 1.000 | 1.000 |
| $\pi d$ | 0.858 | 0.137 | 0.491 | 0.573 | 0.901 | 0.993 | 0.996 |
| $\pi c(0.005)$ | 0.858 | 0.137 | 0.491 | 0.573 | 0.901 | 0.993 | 0.996 |

Ras vs. ERK
|  | EAP | post.sd | 2.50% | 5% | 50% | 95% | 97.50% |
| --- | --- | --- | --- | --- | --- | --- | --- |
| $\Delta\mu$ | **16.382** | 7.041 | 1.36 | 4.493 | 16.695 | 27.203 | 29.469 |
| Size effect | 2.05 | 1.007 | 0.118 | 0.416 | 2.036 | 3.731 | 4.060 |
| Nonoverlap | 0.927 | 0.122 | 0.547 | 0.661 | 0.979 | 1.000 | 1.000 |
| $\pi d$ | 0.881 | 0.126 | 0.533 | 0.616 | 0.925 | 0.996 | 0.998 |
| $\pi c(0.005)$ | 0.881 | 0.126 | 0.533 | 0.616 | 0.925 | 0.996 | 0.998 |

cRaf vs. ERK
|  | EAP | post.sd | 2.50% | 5% | 50% | 95% | 97.50% |
| --- | --- | --- | --- | --- | --- | --- | --- |
| $\Delta\mu$ | **1.680** | 6.527 | -11.45 | -9.028 | 1.753 | 12.104 | 14.377 |
| Size effect | 0.215 | 0.740 | -1.233 | -0.998 | 0.213 | 1.436 | 1.668 |
| Nonoverlap | 0.569 | 0.237 | 0.109 | 0.159 | 0.584 | 0.925 | 0.952 |
| $\pi d$ | 0.554 | 0.184 | 0.192 | 0.240 | 0.560 | 0.845 | 0.881 |
| $\pi c(0.005)$ | 0.553 | 0.184 | 0.192 | 0.240 | 0.560 | 0.845 | 0.881 |

AKT vs. Ras
|  | EAP | post.sd | 2.50% | 5% | 50% | 95% | 97.50% |
| --- | --- | --- | --- | --- | --- | --- | --- |
| $\Delta\mu$ | **-18.970** | 6.94 | -31.053 | -29.014 | -19.571 | -6.964 | -3.532 |
| Size effect | -2.392 | 1.100 | -4.602 | -4.232 | -2.373 | -0.617 | -0.293 |
| Nonoverlap | 0.053 | 0.105 | 0.000 | 0.000 | 0.009 | 0.269 | 0.385 |
| $\pi d$ | 0.09 | 0.113 | 0.001 | 0.001 | 0.047 | 0.331 | 0.418 |
| $\pi c(0.005)$ | 0.09 | 0.113 | 0.001 | 0.001 | 0.047 | 0.331 | 0.418 |

AKT vs. cRaf
|  | EAP | post.sd | 2.50% | 5% | 50% | 95% | 97.50% |
| --- | --- | --- | --- | --- | --- | --- | --- |
| $\Delta\mu$ | **-4.267** | 6.398 | -15.912 | -13.965 | -4.623 | 6.624 | 9.322 |
| Size effect | -0.557 | 0.747 | -2.005 | -1.777 | -0.56 | 0.676 | 0.92 |
| Nonoverlap | 0.327 | 0.222 | 0.022 | 0.038 | 0.288 | 0.751 | 0.821 |
| $\pi d$ | 0.364 | 0.177 | 0.078 | 0.104 | 0.346 | 0.684 | 0.742 |
| $\pi c(0.005)$ | 0.364 | 0.177 | 0.078 | 0.104 | 0.346 | 0.684 | 0.742 |

AKT vs. ERK
|  | EAP | post.sd | 2.50% | 5% | 50% | 95% | 97.50% |
| --- | --- | --- | --- | --- | --- | --- | --- |
| $\Delta\mu$ | **-2.587** | 6.196 | -14.215 | -12.116 | -2.821 | 7.738 | 10.395 |
| Size effect | -0.342 | 0.702 | -1.715 | -1.496 | -0.343 | 0.823 | 1.057 |
| Nonoverlap | 0.389 | 0.223 | 0.043 | 0.067 | 0.366 | 0.795 | 0.855 |
| $\pi d$ | 0.414 | 0.174 | 0.113 | 0.145 | 0.404 | 0.720 | 0.773 |
| $\pi c(0.005)$ | 0.414 | 0.174 | 0.113 | 0.145 | 0.404 | 0.720 | 0.772 |

